# Locus of emotion influences psychophysiological reactions to music

**DOI:** 10.1101/2020.03.28.013359

**Authors:** Julia Merrill, Diana Omigie, Melanie Wald-Fuhrmann

## Abstract

It is now widely accepted that the perception of emotional expression in music can be vastly different from the feelings evoked by it. However, less understood is how the locus of emotion affects the experience of music, that is how the act of perceiving the emotion in music compares with the act of assessing the emotion induced in the listener by the music. In the current study, we compared these two emotion loci based on the psychophysiological response of 40 participants listening to 32 musical excerpts taken from movie soundtracks. Facial electromyography, skin conductance, respiration and heart rate were continuously measured while participants were required to assess either the emotion expressed by, or the emotion they felt in response to the music. Using linear mixed effects models, we found a higher mean response in psychophysiological measures for the “perceived” than the “felt” task. This result suggested that the focus on one’s self distracts from the music, leading to weaker bodily reactions during the “felt” task. In contrast, paying attention to the expression of the music and consequently to changes in timbre, loudness and harmonic progression enhances bodily reactions. This study has methodological implications for emotion induction research using psychophysiology and the conceptualization of emotion loci. Firstly, different tasks can elicit different psychophysiological responses to the same stimulus and secondly, both tasks elicit bodily responses to music. The latter finding questions the possibility of a listener taking on a purely cognitive mode when evaluating emotion expression.

## 1 Introduction

The assumption that the emotion a musical piece expresses is the same as the emotion felt in response to listening to it was long the cornerstone of Western music pedagogy and music-aesthetic discourse (Plato: *Republic*, *Laws*; Aristotle: *Politics;* see (1)). A possible dissociation of the two phenomena has long been theoretically discussed (2–4) and more recently has begun to be empirically examined. In his seminal work, Gabrielsson (5) developed a model of this relationship drawing the distinction between perceived and felt emotions in music. The former is the expression ascribed to a piece of music while the latter is the feeling that it sparks in the listener (also known as “external locus” and “internal locus”, respectively, (6)). The current study aims to identify underlying mechanisms of both loci using continuously measured psychophysiological responses, as these responses have been shown to represent felt affective/emotional states.

It has been theoretically proposed by (5) that these two emotion loci have four distinct relationships: the perceived and the felt emotion could be in a i) positive relationship, ii) negative relationship, iii) in no systematic relationship, iv) no relationship at all. These relationships describe when the individual feels the same emotion as the music is expressing (positive relationship), the individual feels the opposite emotion as the music is conveying (e.g. feeling happy when listening to sad music, negative relationship), feeling a completely unrelated emotion listening to music (e.g. no change in emotional response when listening to music, no systematic relationship), feeling an emotion that cannot be expressed in music (e.g. being moved, no relationship). In terms of their prevalence, these relationships occur in varying frequencies. (7) showed a positive relationship to be by far the most experienced (positive: 61%: negative: 22%: no systematic: 12%; no relationship: 5%). Later, (6) subsumed the four relationships into two: one with both loci in a matched relationship (e.g., a perceived positive emotion leads to a positive emotion in the listener) and another with loci in an unmatched relationship, (e.g., a perceived negative emotion is accompanied by a positive emotion or another feeling which is just not the same as perceived, and therefore including the no (systematic) relationships).

Apart from the prevalence, these and other studies have sought to examine the extent to which felt and perceived emotions show congruency during music listening. Using the dimensional model of emotions, (8) showed that self-reports on felt emotions were stronger than those for perceived emotions when related to pleasure, but were weaker in relation to other affective dimensions. This points to the idea that in the response to different affects or emotions, the loci might enter a different relationship, depending on the music. In a similar vein, (9) examined relationships between felt and perceived emotions in pleasurable and ‘neutral’ music. Relationships between felt and perceived ratings were shown to depend on whether excerpts were self-selected or experimenter-selected: revealing a matched relationship of valence for self-selected music and an unmatched relationship for valence and arousal in experimenter-selected pieces. Specifically, participants reported more neutral experiences for experimenter-selected music, while they reported experiencing pleasure in response to happy and sad musical expression in self-selected pieces. (6) confirmed this result in a meta-analysis by showing that for self-selected (as well as preferred) music, there was a smaller gap across the loci than for experimenter-selected music.

Disregarding specific emotions and other interacting variables, (6) demonstrated, by reviewing 16 studies that used self-reports on the comparison of perceived and felt emotion, that in the majority of cases the emotion felt is congruent with, but often rated lower in intensity than the emotion perceived. Positive expressed emotion was rated higher than positive felt emotion, except in two studies, in which the expressed emotion was higher than the felt emotion (one of them was (8)). “The clearest, new trend, then, that has emerged in the literature since Gabrielsson’s review is that when there is a mismatch between felt and expressed emotion, and when these data are gathered via rating items […], results rarely show that felt emotions are rated as statistically higher than expressed emotions.” (6).

Three explanations for this were pointed out and need to be discussed: Firstly, the instructions given will have an influence on the listener’s differentiation between the loci (as demonstrated by (10)), which means that participants have to understand the difference between perceived and felt emotions in order to be able to fulfill the task. One could also understand the different instructions as different listening modes, i.e. an action-oriented intentional activity of making sense of the world (11), a participant has to adopt in order to fulfill the task (a framwork of listening modes is actually suggested to be able to explain the differences between perceived and felt emotions by (11)). Secondly, because listeners adjust to a certain situation, (social) inhibition processes could occur in response to felt emotions and explain the reduced intensity in laboratory situations (e.g., one would not cry in a laboratory setting in response to sad music; (6)), pointing to the possibility that one might not expect to get extreme bodily responses.

Thirdly, the most plausible explanation for a matched relationship between perceived and felt emotions in music is provided by the mechanism of emotional contagion. A matched relationship could be due to mimickry (insofar as music is able to directly influence the listener in a way that the musical expression is mimicked internally, e.g., peripheral feedback from muscles; (12, 13)) or empathy (14). Unmatched relationships, however, could be explained by idiosyncratic and cultural interferences, which might put a listener in a less ‘susceptible’ state for responding to musical features. They might also be due to the activation of other music-related emotion induction mechanisms, most notably aesthetic emotions, i.e. emotions such as pleasure or boredom that are the consequence of an evaluation of the musical piece (10, 15, 16). Support for this claim could be taken from studies using familiar and highly pleasurable music, revealing higher psychophysiological responses (9). It is of note that this explanation does not account for different degrees of matched/unmatched relationships. One needs to consider though, that music is an artificial and not a natural stimulus with a set of unambiguous expressions, hence, if clues from lyrics or a concrete opera performance are missing, the ‘cause’ for an emotion does not become obvious.

Furthermore, according to the “lense-model” (12), if in the process of a perceived emotion the listener directly decodes emotions through an act of mimicry, psychophysiological reactions should be expected. The latter statement could question a supposed theoretical assumption underlying both loci: While the perceived emotion is understood as a perceptual-cognitive process (such as perception or categorization of an emotional character), the felt emotion is understood as the listener’s emotional response that is feelings reflecting the introspective perception of psychophysiological changes (5, 13, 17–19).

Because most studies on perceived and felt emotions have mainly used self-reports to investigate the relationships, these underlying processes need further investigation with physiological (objective) continuous measures.

### 1.1 Psychophysiological investigations of induced emotions

The study of psychophysiological reactions to music listening is by no means straight forward. Reviews on these relationships have shown diverse results between studies, dependent on tasks, instructions, and music material used (for an overview see (20–22)). Also, the replication of single studies revealed similarities but also differences (23). Studies using psychophysiological measures have investigated music-evoked emotions mainly by asking participants to report on discrete emotions or on affective dimensions (e.g., 24–28). A typical experimental design in this context involves participants evaluating the expression of a musical excerpt by self-reports in one session, and then the same or a new set of participants reporting in another session the emotions felt while psychophysiological measures were recorded. Results often showed matched relationships between both loci, resulting in the conclusion that emotional contagion plays an important role in emotion induction via music (6, 29).

Typical measures for physical arousal comprise heart rate (HR), respiration rate (RR) and skin conductance (SC). For the current approach, studies reporting on effects related to the dimensional model (such as investigating the different reactions to pleasant vs. unpleasant music) are of most interest. Arousing music such as happy, pleasurable and even preferred music (30) results in bodily reaction increases (SC, RR, and HR; 9, 31, 32). Unpleasant music lead to decreases/deceleration in HR (e.g., 26, 33, 34; but see also 35, 36 for no changes), and lead to increases in SC response to fearful music (19). The latter makes a problem obvious in separating valence and arousal in discrete emotions: SC responded stronger to arousing stimuli, such as happy and fearful, than sad and peaceful music (19, 32). Also, an assumption that seems to exist is that sadness in music is unpleasant, which is not always true as sad music can be pleasurable (16, 37). Therefore, musical excerpts used in the current study were chosen to represent the poles of the affective dimensions, namely negative and positive valence, and high and low energy and high and low tension.

Facial electromyography (EMG) is also typically recorded alongside psychophysiology to measure experiences. As measure of muscle activity, it is usually recorded through the zygomaticus major muscle (38), associated with smiling and therefore positive valence, and the corrugator supercilii muscle, associated with frowning and therefore negative valence (for pleasant and unpleasant stimuli, see 39, 40). Accordingly, positively valenced music has been associated with increases in zygomaticus activity while negatively valenced music has been associated with an increase in corrugator activity (in combination with movies, see 41, 42). This clear-cut pattern, however, has not consistently been reported in music studies (e.g., 27, 32). In contrast to arousal measures, which are considered measures of psychophysiology, EMG is considered a measure of behavior (43). This points to the idea that EMG could be understood to also indicate aesthetic judgements, which has been shown with higher corrugator activity for disliked dance clips and higher zygomaticus activity for liking (44). This could explain some of the previous heterogeneous results (e.g., zygomaticus activity could be the result of a positive aesthetic evaluation of a negatively valenced stimulus).

Only one study gives rise to the assumption that differences relative to the task given to participants could be visible in the autonomic nervous system. (45) explored the effect of different task instructions on psychophysiological responses to opera music. They observed that instructions to either adopt an ‘objective’ or an empathic perspective while listening to and watching a video of a ‘happy’ aria and a ‘sad’ song led to significant differences in psychophysiological responses. Participants additionally reported on how strongly they felt empathy with the singer and the story the soprano was experiencing. Physiological reactions reflected the emotions experienced by the character in the ‘high empathy’ condition differently than in the ‘low empathy’ condition. However, this study used video, instead of music alone, it is difficult to draw coherenct conclusions from these psychological difference as it has been shown that psychophysiology responds differently to emotional performance between audio, visual and audio-visual performance (46, 47). Therefore, more research needs to be conducted observing psychophysiology in different conditions for music listening only.

Another study investigated how the central nervous system reacts to differences between the two emotion loci using functional magnetic resonance imaging (fMRI). In the felt emotion task, participants were asked whether they experienced the emotion in a musical excerpt and in the perceived emotion task, if the music expresses the respective emotion, which was happy and sad (48). This failed to show a significant difference between both tasks. However, contrasts to baseline (listening without task) showed greater activation in bilateral inferior frontal gyri (left Brodmann area (BA) 47, right BA 45) during the perceived task, which, according to the author’s interpretation, reflects the assessment of musical expression, and in the precuneus (BA 7) during the felt task, possibly reflecting self-representational processes of assessing own emotional responses. These results seem to give support to the above stated difference of self-evaluation on the one side and detecting musical expression on the other.

### 1.2 The present investigation

Relations between psychophysiological measures and emotion perception have been investigated under different premises, methods as well as concepts and terminology (see for example the discussions on aesthetic emotions and utilitarian/real emotions, (10, 49–51)). The inherent problem lies in the multidimensionality of both affective processing and expression in music. A first step to reducing this complexity is to test the two emotion loci of perceived and felt emotions with simpler tasks (and only audio) while measuring psychophysiology. Hence, the current study compares the psychophysiological correlates of the act of perceiving musical expression with those of the act of assessing feelings induced by music listening.

Stimuli and rating scales corresponded to a study by Eerola and Vuoskoski (52). For the perceived task, we asked participants to choose one adjective out of four, representing two dimensions: ‘positive, negative, tense, relaxed’. For the felt task, we asked our participants to report on the intensity of their evoked feelings only, to check that the stimuli were capable of inducing emotions of different intensities. After the study, participants reported on the strategies used to fulfill the task in a qualitative methods approach to gain further insights into the processes of performing the two tasks and to interpret the responses from the psychophysiological measures.

### 1.3 Hypotheses

Firstly, with regard to physiological response as a function of affective dimensions, we hypothesized that high arousal would result in greater skin conductance, respiration rate and heart rate than low arousal clips (as already shown in previous studies, see above). Under the assumption of viewing facial EMG as a measure of behavior, we also predicted that positive valence would result in greater zygomaticus activity than negative valence and that negative valence would result in greater corrugator activity than positive valence reports.

Secondly, we hypothesized that the locus of emotion would influence the amplitude of physiological responses: If the act of introspecting on one’s emotional state boosts emotional response, we would expect to see a greater physiological response for the felt emotion task in which the emotion locus is internal. If the act of perceiving includes an analytical assessment without emotional involvement, we can expect suppressed and reduced responses, if it necessarily involves simulation of the emotion, we can expect that the physiological response would be at least as large as for the felt task. Finally, if, however, the act of focusing on the internal emotion locus distracts from the music itself, we should expect to see larger physiological responses during the perceived task.

## 2 Methods

### 2.1 Ethics statement

All experimental procedures were ethically approved by the Ethics Council of the Max Planck Society, and were undertaken with written informed consent of each participant.

### 2.2 Participants

40 participants (mean age = 26.48 years, SD = 7.77; 12 males) were tested. Participants showed various musical backgrounds (mean musical instrument playing = 7.28 years (SD = 6.58), range 0-20 years).

### 2.3 Materials

Stimuli were taken from movie soundtracks, whose musical and stylistic properties as well as the film context they originated from, were considered suitable for the differentiation between felt and perceived emotions. 32 musical excerpts were selected on the basis of the ratings reported in Eerola and Vuoskoski (52). They were selected based on the emotion groupings (Table 1) with highly and moderately representative examples of the poles of the respective affective dimension to keep the participants’ attention focused on the musical expression in the perceived task: positive (*N* = 8) and negative rated valence (*N* = 8), low (*N* = 4) and high tension (*N* = 4) as well as low (*N* = 4) and high energy (*N* = 4). The original 15 second pieces reported in the study were extended to approximately 30 second excerpts and maintained a constant musical expression over their duration (see also (53)) to account for the rather slow psychophysiological measures.

**Table 1.** List of stimuli. L = looped stimuli. The last five columns refer to the final stimulus set ‘2’ from (52). The chosen stimuli form the ‘Emotion Grouping’ consisted of highly (HIGH) and moderately (MOD) representative examples of the poles of the respective emotion dimensions.

| Excerpts in current study |  |  |  | Stimulus set '2' from Eerola and Vuoskoski (2011) |  |  |  |  |
| --- | --- | --- | --- | --- | --- | --- | --- | --- |
| Album Name | Track | Start | End | Index Reference | Ratings: Valence | Energy | Tension | Emotion Grouping |
| Batman | 4 | 02:31 | 02:48L | 79 | 7.25 | 8.12 | 5.03 | ENERGY POS MOD |
| Batman | 9 | 00:56 | 01:23 | 63 | 4.76 | 3.96 | 6.04 | VALENCE NEG MOD |
| Big Fish | 8 | 00:11 | 00:42 | 110 | 6.75 | 3.90 | 2.82 | TENSION NEG HIGH |
| Big Fish | 15 | 00:18 | 00:49 | 65 | 4.07 | 5.04 | 6.93 | VALENCE NEG MOD |
| Big Fish | 15 | 00:54 | 01:26 | 82 | 5.24 | 3.70 | 4.79 | ENERGY NEG MOD |
| Blanc | 12 | 00:39 | 01:09 | 52 | 6.82 | 4.97 | 3.46 | VALENCE POS HIGH |
| Blanc | 10 | 00:12 | 00:42 | 88 | 5.03 | 2.96 | 4.10 | ENERGY NEG HIGH |
| Cape Fear | 2 | 01:13 | 01:44 | 100 | 3.22 | 5.09 | 7.43 | TENSION POS MOD |
| Dances with Wolves | 10 | 00:18 | 00:46 | 55 | 7.04 | 7.27 | 4.66 | VALENCE POS HIGH |
| Dracula | 5 | 00:11 | 00:44 | 99 | 4.16 | 5.22 | 6.84 | TENSION POS MOD |
| Gladiator | 17 | 00:00 | 00:29 | 53 | 7.07 | 6.76 | 4.55 | VALENCE POS HIGH |
| Hellraiser | 5 | 00:00 | 00:30 | 69 | 2.36 | 7.09 | 8.24 | VALENCE NEG HIGH |
| Juha | 2 | 01:55 | 02:26 | 101 | 7.57 | 4.60 | 2.46 | TENSION NEG MOD |
| Juha | 10 | 00:07 | 00:40 | 51 | 7.30 | 5.00 | 3.18 | VALENCE POS HIGH |
| Juha | 16 | 00:00 | 00:31 | 81 | 5.60 | 3.07 | 4.01 | ENERGY NEG MOD |
| Juha | 18 | 02:28 | 03:00 | 61 | 5.78 | 5.97 | 5.99 | VALENCE NEG MOD |
| Lethal weapon 3 | 7 | 00:00 | 00:29 | 67 | 4.72 | 4.72 | 6.67 | VALENCE NEG HIGH |
| Man of Galilee CD1 | 2 | 00:09 | 00:42 | 56 | 6.67 | 6.75 | 5.16 | VALENCE POS MOD |
| Man of Galilee CD1 | 2 | 03:02 | 03:18L | 72 | 7.45 | 8.39 | 5.61 | ENERGY POS HIGH |
| Oliver Twist | 7 | 00:53 | 01:20 | 80 | 5.88 | 6.78 | 6.10 | ENERGY POS MOD |
| Pride & Prejudice | 4 | 00:02 | 00:33 | 105 | 5.84 | 5.90 | 5.67 | VALENCE POS MOD |
| Road To Perdition | 6 | 00:18 | 00:49 | 68 | 8.01 | 8.27 | 4.16 | TENSION NEG MOD |
| Running Scared | 15 | 01:44 | 02:16 | 86 | 2.51 | 6.03 | 8.01 | VALENCE NEG HIGH |
| Shakespeare In Love | 11 | 00:20 | 00:51 | 62 | 6.57 | 7.55 | 5.51 | VALENCE POS MOD |
| Shakespeare In Love | 21 | 00:00 | 00:29 | 57 | 4.28 | 3.88 | 5.84 | ENERGY NEG HIGH |
| The Alien Trilogy | 11 | 02:05 | 02:35 | 91 | 3.48 | 5.58 | 7.37 | VALENCE NEG MOD |
| The English Patient | 8 | 01:28 | 01:59 | 66 | 3.69 | 7.09 | 7.66 | TENSION POS HIGH |
| The Fifth Element | 13 | 00:16 | 00:48 | 92 | 2.58 | 6.85 | 8.16 | VALENCE NEG HIGH |
| The Godfather Part III | 5 | 01:11 | 01:41 | 107 | 2.99 | 6.99 | 8.12 | TENSION POS HIGH |
| The Untouchables | 6 | 01:38 | 02:08 | 71 | 7.09 | 4.21 | 2.75 | TENSION NEG HIGH |
| Vertigo OST | 6 | 02:02 | 02:33 | 58 | 7.45 | 7.64 | 4.27 | ENERGY POS HIGH |
| Outbreak | 6 | 00:15 | 00:45 | 60 | 6.03 | 3.82 | 4.31 | VALENCE POS MOD |

### 2.4 Procedure

Participants were informed about the study and carefully instructed regarding the meanings of and difference between the concept of perceived and felt emotions. They were then prepared for the recordings of psychophysiological measurements, which included the measurement of the zygomaticus major muscle (EMGZ) and corrugator supercilii muscle (EMGC) with Ag/AgCl electrodes, respiration rate (RR) with a custom prepared respiration belt wrapped around the upper rib cage (level with sternum), skin conductance (SC) with electrodes attached to the index and middle fingers of the left hand (which was instructed to remain still throughout the experiment), as well as a plethysmograph clip used to measure the blood volume pulse (BVP), from which heart rate (HR) could be inferred.

For facial EMG, electrodes were placed on the left side of the face according to (38) and (54) guidelines. Two electrodes were placed over the eyebrow to measure the activity of the corrugator muscle, while two others were placed on the cheek to measure the zygomaticus muscle activity. The ground electrode was placed in the middle of the forehead. Impedance of facial EMG electrodes was kept below 5 kΩ. Electrodes were connected to the amplifier with a 10 s low cutoff for EMG and a DC-cutoff for arousal measures and a 250 Hz high cutoff (to reduce electrical interference). Activity was sampled at a 500 Hz rate.

Participants were seated in front of a monitor and listened to the music over loudspeakers (Neumann KH 120 A). Loudness was adjusted to a comfortable level for each participant. Participants were instructed to make all ratings with a computer mouse using their right hand, and to avoid moving their left hand during recordings. A set of six training trials was performed in order to give the participants an impression of the study and to familiarize them with the two different tasks.

Each trial commenced with a prompt on the screen indicating one of the two tasks, either perceived (German ‘Ausdruck’ = expression) or felt (German ‘Gefühl’ = feeling) for 4 seconds. Participants were instructed to carry out the two tasks as follows: “One task asks for the expression you perceive in the music. To describe this, please select one of the four given adjectives: ‘positive, negative, tense, relaxed’. It is possible that more than one could fit; please choose the one, which in your opinion fits the best; there is no right or wrong answer. The other task asks for the intensity of feeling that the music evokes in you. Focus on yourself and observe your own reaction. The music can touch you a little or greatly. Please indicate this on a scale from 1 (little) to 5 (strong). Here, no descrete description of your feeling is needed.” After the music ended, the rating scale for the respective tasks appeared. The task to choose an adjective was only to help participants to focus on the task because for the current study, it was only of interest to continuously record the process of perceiving or feeling an emotion in/through music.

In combination with psychophysiological measures, we opted for mixing the tasks as well as musical pieces in one session to maintain high attention in participants’ (6, 10). Trials were grouped in four blocks with the opportunity for breaks in between them. Each block started with a 30 second rest period before the presentation of a fixed and pre-set number of musical excerpts in randomized order. The two tasks also occurred randomly within a block but were balanced in numbers. Participants started with different blocks to assure every stimulus was presented in the context of one task in a different order. Over the course of a recording session, each excerpt was played twice, once in the context of each task. Following the approximately 50 minute recording session, participants filled out a debriefing questionnaire in which they were asked about the strategies used to fulfill the two tasks.

### 2.5 Analysis

All signal processing was carried out using custom scripts written in MATLAB 2017b (55). To obtain a continuous measure of facial muscle activity over time, the zygomaticus and corrugator facial muscle signals were band-pass filtered between 20 and 249 Hz, and the absolute value of the Hilbert transform of the filtered signal was extracted and then convolved using the *conv* function in MATLAB as recommended (http://www.fieldtriptoolbox.org/documentation; 56). Heart and breathing rate over time were estimated over several steps. In the first step, the raw respiration and BVP signal were low pass (40 Hz) and high pass filtered (0.05 Hz) to remove body and hand movement artifacts, as well as drifts that can affect the ability to identify peaks reliably. Next, peaks in the signal were identified using functions that identified local minima and maxima with visual inspection. Finally, rate for each signal was estimated by taking a differential of the timings of maxima *(1./diff(timing of peaks)),* and the resulting rate time-series were interpolated to a regular sampling frequency.

To obtain a time-series of the phasic component of the skin conductance data (SC) for each piece, functions from the *Ledalab* toolbox V.3.4.9 were used (57). Note that no threshold was used for estimation of SC but rather, as recommended (57), a time integration of the continuous measure of phasic activity (average value) was taken as a direct indicator of event-related sympathetic activity.

Finally, to reduce the inter-individual variance of absolute amplitudes for the physiological data, activity for all physiological measures during the music listening tasks was normalized across all trials within a participant.

For statistical analysis, the mean response of the first and the second half of the musical pieces (i.e. 15 second intervals) was calculated to account for changes in physiological measures during the duration of musical pieces (cf. (58)).

Analysis was based on the expressiveness ratings provided by Eerola and Vuoskoski (52) which were collected from a large group of participants (*N*=116). It is of note that a comparison with the ‘perceived’ emotion ratings from our own participants was not planned, and because of our design, their answer only represented one dimension, either valence or arousal, which did not allow for further comparisons. For statistical analyses, the independent variables of affective dimension were grouped into three categories based on the rating scale from 1-9 (52), i.e., for Energy and Tension, 1-3 = low, 4-6 = medium, 7-9 = high, and for Valence, 1-3 = negative, 4-6 = neutral, 7-9 = positive.

### 2.6 Statistical analysis

Statistical analyses were carried out using *R* version 3.5.1 (59). Supplementary material can be found here: https://edmond.mpdl.mpg.de, collection Locus of Emotion in Music.

#### 2.6.1 Effects of emotions and tasks on psychophysiological measures

The *lme4* package (60) was used to perform linear mixed effects models (LMM) and examine how the different physiological variables responded to the experimental manipulations. To confirm the basic assumptions underlying LMMs, we visually inspected residual plots, quantile-quantile plots, and grouped boxplots of residuals, representing linearity, normality, and homoscedasticity.

Each psychophysiological measure was separately analyzed. We took as fixed effects in the linear mixed models one of the affective dimensions with three levels each, that is: Valence (negative, neutral, positive), Energy (high, medium, low), or Tension (high, medium, low), along with Task (felt, perceived). As we had no hypotheses regarding relationships between the affective dimensions, we did not include them into one model but fitted three separate ones.

Models were fitted with random intercepts of participant, item (i.e., musical piece) and interval (first and second half) to account for these influences. Random slopes were not included because no variance between participants with regard to Emotion or Task had to be accounted for as we used external ratings and items were the same between conditions.

*F*- and *p*-values were obtained by the *anova* function from the *car* package (61) and confidence intervals were calculated using the *confint* function from the *lme4* package. Two (Pseudo-)R^2^ values were calculated with the *MuMin* package (62), which is designed for general and linear mixed effects models. The *r.squaredGLMM* function produces two coefficients of determination, one represents the variance explained by the fixed effects (‘marginal’ R^2^(m)_GLMM_), the other the variance explained by the entire model (‘conditional’ R^2^(c)_GLMM_), including fixed and random effects (Table 2).

**Table 2.**
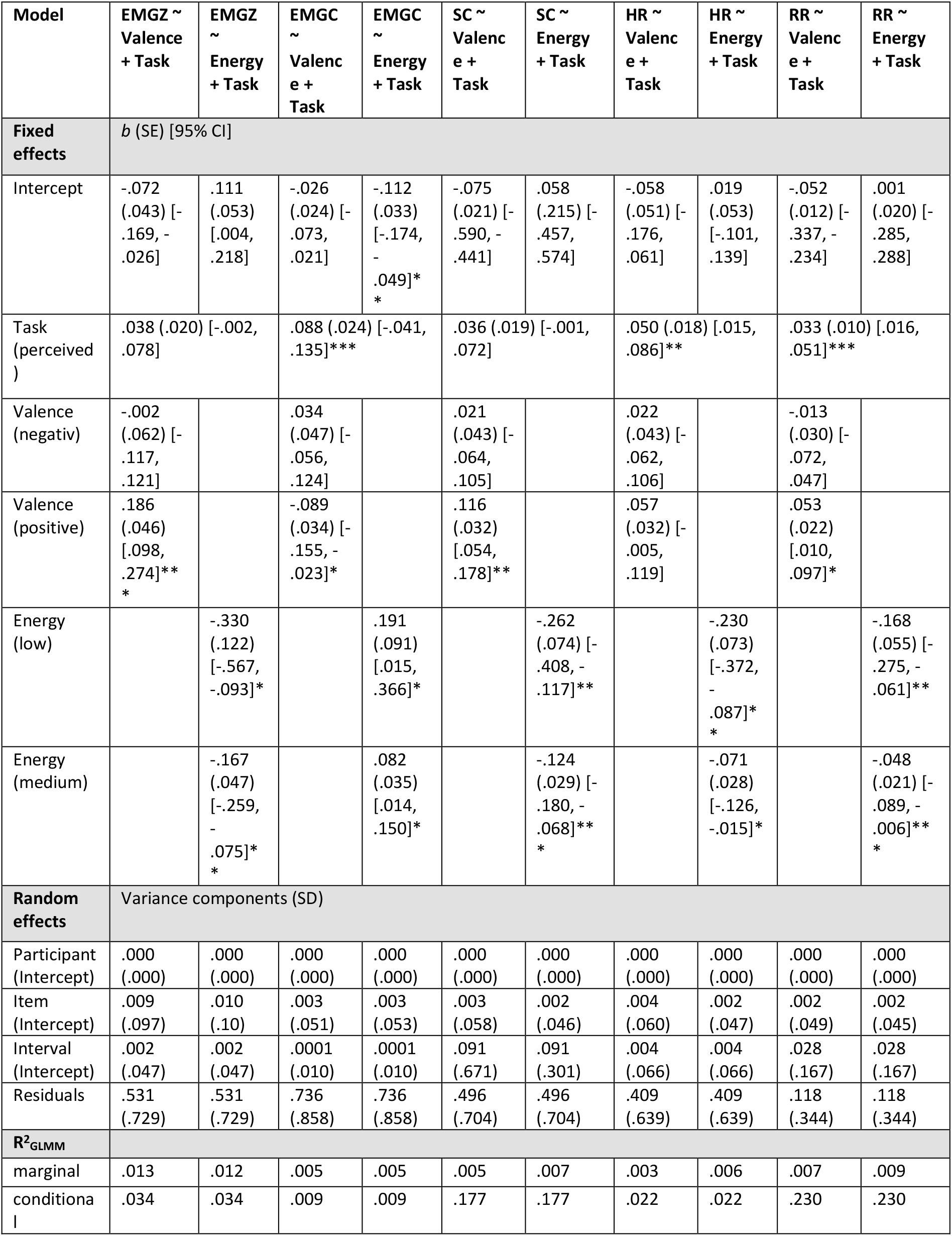
Fixed effects and random effects for models with Valence, Energy and Task. For fixed effects, the beta estimate (b) with Standard error (SE) and 95% confidence intervals (CI) are reported, for random intercepts, the variance components and standard deviation (SD). For each model, two types of R^2^_GLMM_ are reported, marginal (m) and conditional (c) (see text for more information). Note that fixed effects are interpreted based on the *F*-tests reported in the text. Significance codes: *p* < .001 ‘***’ *p* < .01 ‘**’ *p* < .05 ‘*’.

#### 2.6.2 Participants’ ratings and self-reports

To evaluate the relationships between loci, we investigated whether the felt intensity ratings matched the perceived emotion ratings (as rated by the participants in the study by (52)) and whether they were reflected in the psychophysiological measures. To this end, two additional analyses were carried out. Firstly, the participants’ self-reported intensity ratings were correlated (Pearson) with the perceived emotion ratings by Eerola and Vuoskoski (52). Secondly, the intensity ratings were compared to the psychophysiological measures using linear mixed effects models. The models were fitted with z-transformed intensity ratings taken as fixed effects.

From a debriefing form with open questions the participants filled out after the study, we derived categories on the strategies the participants used to solve the task.

**Figure 1.**
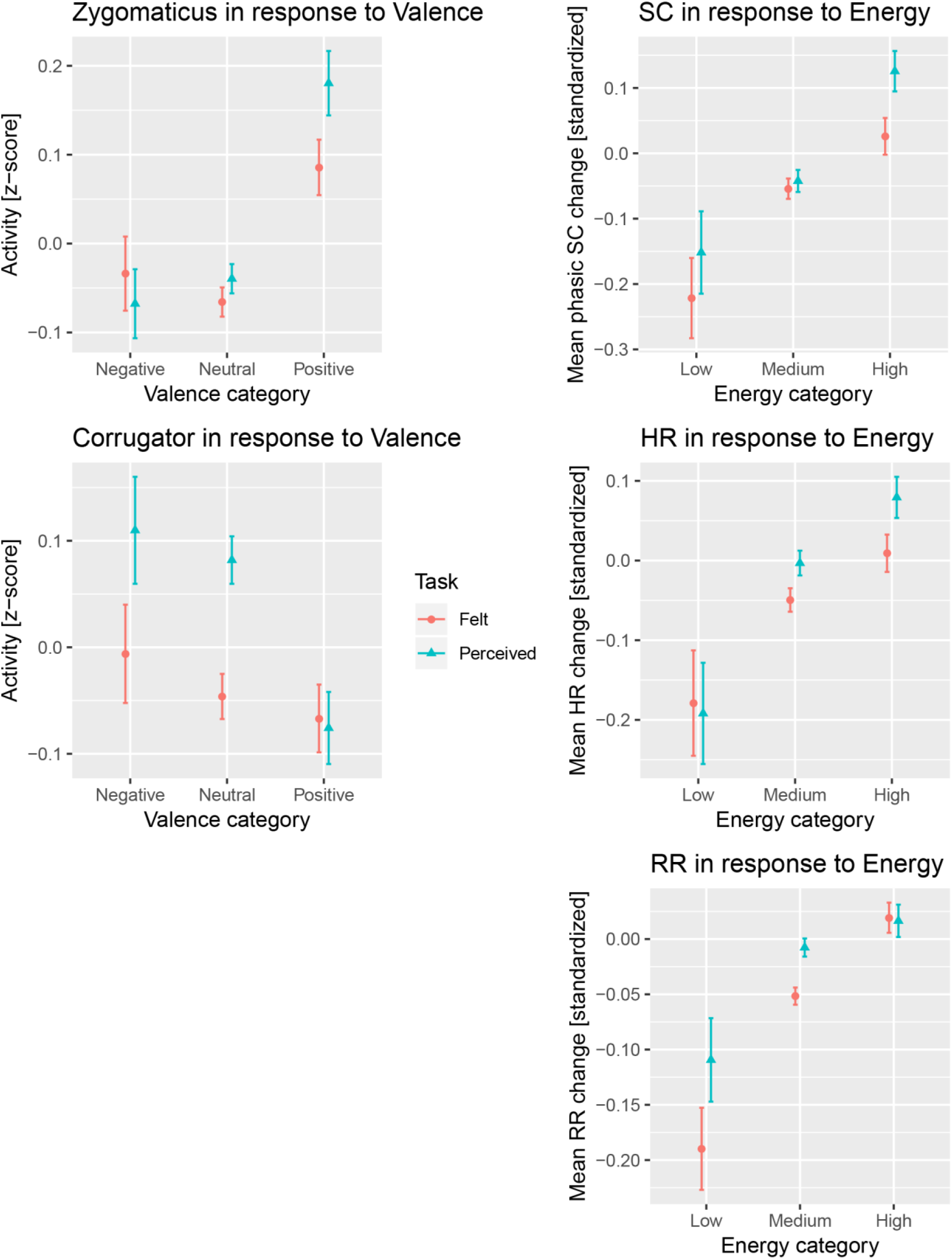
The mean activity of zygomaticus and corrugator muscles in response to Valence (left column) and from skin conductance, heart and respiration rate in response to Energy (right column). On the x-axis the respective categories are depicted and on the y-axis the standardized mean activity/response.

## 3 Results

### 3.1 Zygomaticus muscle activity

Valence affected EMGZ activity (*F*(2,29) = 8.781, *p* = .001), increasing its activity from neutral to positive valence by 0.186 ± 0.046 (Estimate ± standard error, SE) and decreasing it slightly from neutral to negative by 0.002 ± 0.020 SE (for details, see Table 2). Energy affected EMGZ (*F*(2,29) = 7.820, *p* = .002), increasing its activity from low to medium to high energy (for parameter estimates, see Table 2). Tension did not show a significant effect (*F*(2,29) = 0.896, *p* = .419). Task only marginally affected EMGZ (*F*(1,5086) = 3.477, *p* = .062), with slightly higher activity for felt than perceived by 0.038 ± 0.020 SE.

### 3.2 Corrugator muscle activity

Valence affected EMGC activity (*F*(2,29) = 4.320, *p* = .023), increasing its activity from positive to neutral to negative valence. Energy affected EMGC (*F*(2,29) = 3.774, *p* = .035), increasing its activity from high to medium to low. Tension did not affect EMGC (*F*(2,29) = 2.162, *p* = .133). Task affected EMGC (*F*(1,5086) = 13.464, *p* < .001), with higher activity for felt than perceived. Note that the R^2^_GLMM_ values for these models are the smallest compared to other measures in the current study, i.e., the effects of Valence, Energy and Task on EMGC have to be considered small.

### 3.3 Skin Conductance response

Energy affected the SC response (*F*(2,29) = 12.165, *p* < .001), increasing its response from low to medium to high energy. Tension did not affect SC (*F*(2,29) = .475, *p* = .627). Valence affected SC (*F*(2,29) = 6.760, *p* = .004), increasing its response from middle to positive valence and slightly from middle to negative valence. Task affected SC only marginally (*F*(1,5086) = 3.602, *p* = .058), with higher activity for felt than perceived.

### 3.4 Heart Rate response

Energy affected HR (*F*(2,29) = 6.317, *p* = .005), increasing its response from low to medium to high energy. Neither Valence (*F*(2,29) = 1.617, *p* = .216) nor Tension (*F*(2,29) = .294, *p* = .748) affected HR response. Task affected HR (*F*(1,5086) = 8.015, *p* = .005), with higher response for felt than perceived.

### 3.5 Respiration Rate response

Energy affected RR (*F*(2,29) = 5.603, *p* = .009), increasing its response from low to medium to high energy. Valence affected RR (*F*(2,29) = 3.378, *p* = .048), increasing its response from neutral to positive and slightly from neutral to negative valence. Tension did not affect RR (*F*(2,29) = 2.070, *p* = .144). Task affected RR (*F*(1,5086) = 13.722, *p* < .001), with higher response for felt than perceived.

### 3.6 Felt-intensity ratings

The participants’ self-reported intensity ratings were correlated with the perceived emotion ratings by Eerola and Vuoskoski (52) and revealed significant positive correlations with Energy (Pearson *r* = .43, *p* < .05) and Tension (*r* = .42, *p* < .05), showing that the response was related to perceived arousal as intended, but not with Valence (*r* = −.30, *p* = .10) (Figure 2).

**Figure 2.**
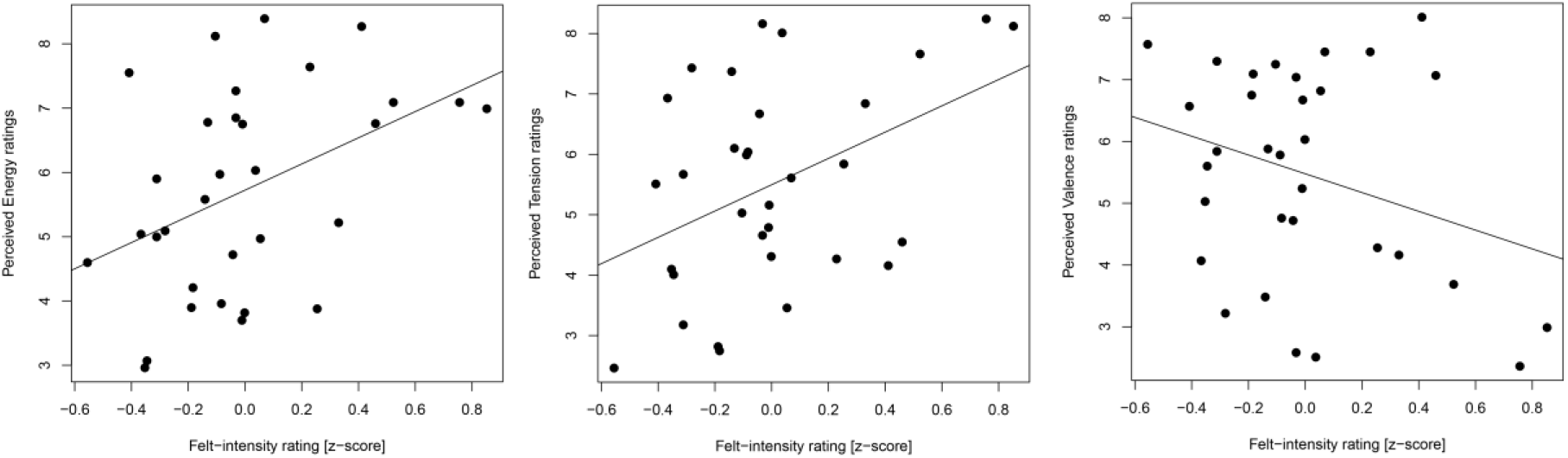
Scatterplots of felt-intensity ratings (z-score) during the felt task (x-axis) and perceived emotion ratings from Eerola and Vuoskoski (52) (y-axis) for all 32 musical pieces.

Relations between the felt-intensity ratings and the psychophysiological measures were investigated with linear mixed effect models, which were fitted as described above and with z-transformed intensity ratings as fixed effects. EMGZ was affected by the felt-intensity ratings (Chi^2^(1) = 12.194, *p* < .001), increasing its activity with higher felt intensity by 0.051 ± 0.014 SEs z-scores. RR was also affected (Chi^2^(1) = 10.347, *p* = .001), increasing its response by 0.022 ± 0.007 SEs. Other measures were not affected by the felt-intensity ratings: EMGC (Chi^2^(1) = 1.917, *p* = .166), SC (Chi^2^(1) = 2.915, *p* = .088) slightly increased its response by 0.023 ± 0.014 SEs, HR (Chi^2^(1) = 2.738, *p* = .098) slightly decreased its response by 0.021 ± 0.013 SEs.

### 3.7 Strategies for solving the tasks

In order to be able to interpret the continuous psychophysiological responses, participants were asked in a debriefing session to describe the strategy they used to carry out the tasks. Participants reported paying attention to one or more of the typical cues of expressivity in Western music (such as tempo, harmonic progression, loudness, timbre, melody or musical structure) during the perceived emotion task. They also reported trying to imagine a real-life situation or movie scene to which the piece could serve as background music or musical illustration.

In contrast, the focus of the felt task was directed inward in line with the instructions. Participants described their strategy with a German metaphor for introspection (to “listen into themselves”) and reported three kinds of cues to evaluate their feelings: i) bodily reactions (like goosebumps, chills, heartbeat), ii) being moved, touched or carried away by the music, and iii) aesthetic judgments about the music (liking and interest).

## 4 Discussion

The current study aimed to investigate potential differences in psychophysiological measures during music listening between two emotion loci: an external and an internal locus, or, in other words, the perceived emotion and the felt emotion. Our results showed stronger bodily reactions when participants focused on an external locus, experienced as focusing on the music, than when focusing on an internal locus, i.e. directed inward in order to assess the emotion induced by music. Our findings have implications for the understanding of physical reactions to the emotions conveyed and elicited while listening to music, and more generally, for dissecting responses to objective acoustical properties from the assigned musical and aesthetic qualities, their ‘content’ (such as emotional expressivity) and the listening mode participants can adopt.

### 4.1 Effects of Task

Results revealed a higher mean response for the perceived emotions task in psychophysiological measures. The effect became significant in corrugator muscle activity, HR and RR response. For the zygomaticus muscle and skin conductance responses, the effect marginally failed to reach significance. This result is in line with previous studies (6–8, 48, 63) that have shown participants’ self-reports to be higher for the perceived emotion in music than the felt emotion. One other study has investigated the effect of different instructions on psychophysiological responses to the same music performance (45). While this study is only partly comparable to our study since it did not include a task in which participants had to focus solely on themselves, the results point in the same direction: When feeling with the performer/music (‘high empathy’) in contrast to ‘not getting caught up’ (‘low empathy’), bodily reactions were more influenced by the expression of the piece (SC level decreased supposedly because of the nostalgic/sad expression and RR increased because of the positive expression). One can conclude that focusing on the main ‘attraction’ in the music (performer, instrument, melody etc.) leads to enhanced psychophysiological reactions. Thus, we conclude from the current study that the focus on one’s self distracts from the music itself (in effect operating as a distractor (task)) and leads to less responsive bodily reactions. In contrast, paying attention to the expression of the music and consequently to tempo, timbre, loudness, harmony (all strong predictors of changes in autonomic response), enhances bodily reactions. This interpretation of the source of the enhanced physiological responses is reflected in the participants’ self-reports obtained in the debriefing. During the perception task, they reported having mostly paid attention to musical features and imagined situations that matched with the musical expression. The latter strategy in particular could be seen as an effective way to induce emotions via empathy (e.g., (45)). During the felt emotion task, on the other hand, they reported having directed their attention to their own (bodily) reactions and aesthetic feelings such as being moved or touched by the music as well as aesthetic judgments such as liking.

Methodologically speaking, the instructions given to the participants for the perceived task might be interpreted as being more effective in inducing emotions in the listener given their inherent invitation to focus on the musical details and to empathize with the perceived emotional expression - at least for musical stimuli of such short duration. Therefore, our results contribute to research on the cognitive processes behind the perception of musical expression, i.e., whether musical emotions should be conceptualized as cue decoding processes or as the subjective interpretation of physiological activation or feelings (e.g., 10, 12, 13). We suggest that, even though participants rely on external information during perception tasks, a purely cognitive mode of emotion decoding without any sort of mimicry or empathy may not be possible in such tasks. We suggest that directing attention inward (as in the felt task) may be a problematic emotion induction task as music related responses are potentially reduced or masked. Here, an instruction asking participants to open themselves up to the music and let it touch them might have led to different results. It is worth pointing out, however, that focusing on one’s own felt emotion might actually lead to higher psychophysiological responses than perceived emotions when strong personal attitudes influence bodily reactions such as in the specific case of highly pleasurable and self-selected music (8, 9). As the music presented to the participants was not self-selected, this might explain why only in some cases the felt emotions was rated higher than the expressed emotion in music in these particular studies as pointed out by (6).

Even though a direct investigation of the influence of specific emotions on the two tasks was not intended with the current study, a matched relationship between loci (in the sense of a ‘positive’ relationship, (5)) is suggested by the participants’ felt intensity ratings that reflect expressed energy and tension. In the bodily response, though, a slight relationship could be seen between high intensity ratings on the one hand and zygomaticus activity and respiration rate on the other, suggesting that an intense felt emotion was positively valenced, regardless of the valence of the musical expression. Therefore, self-reports on felt intensity might be based on aesthetic judgments as also suggested by the participants’ reported strategies on how to solve the task (such as being moved or experiencing pleasant feelings etc.). Hence, a straightforward interpretation of bodily responses via emotional contagion (which would explain a matched relationship) needs to be re-evaluated, because aesthetic judgments have to be taken into account as an emotion (co-)induction mechanism (as has previously been argued by (64)).

These findings may inspire future research on the effects listening modes have on the experience of music (cf. (11)). This would account for one of the main differences aesthetic theories claim to exist between artworks and stimuli from the every-day world, that is that artworks do not only or even primarily consist of their physical properties but are to various degrees co-created by the perceiver in the act of perceiving (see (65) for the theory of the open work). Accordingly, co-creation strategies should be expected to change the object perceived and thus not only higher-order but also lower-order responses to it.

### 4.2 Effects of Valence and Arousal

Our results showed patterns that were largely in line with previous studies. With regard to valence, the zygomaticus muscle was greater for music with a positive expression and the corrugator muscle in response to negative expression (e.g., 26, 42). As the valence effect of the corrugator muscle was not particularly large and other studies showed heterogeneous results with regard to facial muscle reactivity (e.g., 27, 32), one should also consider that music in general is intended to elicit pleasure in its listeners even in response to sad or otherwise negatively valenced music. This has been thoroughly discussed in the context of mixed (37) and aesthetic emotions (16) and will need further investigations with regard to psychophysiological responses.

Reports of perceived high energy correlated, as hypothesized, with SC, HR and RR (increased responses to higher energy), but also with facial muscle activity (increase in EMGZ and decrease of EMGC with higher energy). Previous studies used mainly stimuli representing discrete emotions, showing that arousing music such as highly pleasurable music or happy and fearful music has an effect on arousal measures (9, 19, 31, 32). This approach though, does not always allow to separate arousal from valence effects. Differentiating into affective dimensions such as valence, energy and tension, we showed that SC and RR (albeit the latter with a small effect) were higher in response to positive than negative valence; HR was not affected by valence. With regard to valence measures, zygomaticus activity was higher in response to high energy music and corrugator activity was lower. While further studies are needed to investigate the role and interplay of psychophysiological measures in reported valence and arousal, one might suggest from the current study that valence effects in music might be difficult to be separated from arousal effects because a perceived degree of arousal can have a felt affective valence and the other way round.

Finally, perceived high tension did not correlate with any of the measures we recorded. Eerola and Vuoskoski (52) already noted that two dimensions (of valence, tension and energy) were a good model fit in explaining perceived emotions in music, also showing high correlations between energy and tension, supporting the idea that one of the dimensions is redundant in reflecting arousal in music (see also (66) for further investigations of tension and energy in physiological response to specific ‘beautiful’ musical passages). Based on psychophysiology, our study suggests tension to be the one to eliminate. One could also conclude that behaviorally rated tension in music may not be well reflected in bodily reactions, because of its interplay with relaxation and its major component of uncertainty (67), which needs further investigation.

### 4.3 Conclusion

The current study investigated differences between the external and internal emotion locus in music with regard to psychophysiological measures that index subjective experiences. While earlier studies compared ratings of emotions perceived to be expressed by musical pieces (exernal locus) with self-reports on felt emotions induced by these pieces (internal locus) and mostly found the perceived ratings to be stronger than the felt emotions, our study compared the effect of two instructions intended to induce a listening mode either focusing on emotion perception or on induced emotion on psychophysiological responses. Similarly to the behavioral studies, but still surprising in comparison with other psychophysiological studies inducing strong emotions such as pleasure, the physiological reaction to the perceived emotion task was stronger than to the felt task. While during the perceived task, bodily reactions likely evolve due to a focus on acoustical and musical properties and musical imagery (as seen in the reported strategies by the participants), response patterns during the felt task are likely weaker due to introspection and hence distraction from the music itself. With regard to research on psychophysiological markers of music listening and facets of the aesthetic experience of music, our study also showed that one and the same musical stimulus can result in different psychophysiological responses depending on the listening mode a person adopts, due to differing instructions. Even though sympathetic activation is generally thought to be an automatic and comparatively low-level process caused by basic acoustic properties of musical pieces, top-down processes come into play. Evoked feelings as well as emotion decoding processes in music involve bodily reactions.

## 5 Acknowledgement

We thank Sandro Wiesmann, Freya Materne and Elena Felker for help with data collection, Cornelius Abel for laboratory support and Pauline Larrouy-Maestri, Anna Czepiel and Christoph Seibert for comments on an earlier version of the manuscript.

## 7 Figure Supplement

Figure S1. Plots for all measures and affective dimensions. The mean activity/response of zygomaticus major muscle (row 1), corrugator supercilii muscle (row 2), skin conductance (SC, row 3), heart rate (HR, row 4), respiration rate (RR, row 5) in response to valence (column 1), energy (column 2) and tension (column 3). On the x-axis the respective categories are depicted and on the y-axis the standardized mean activity/response.

